# Evidence of attack deflection suggests adaptive evolution of wing tails in butterflies

**DOI:** 10.1101/2022.04.05.487106

**Authors:** Ariane Chotard, Joséphine Ledamoisel, Thierry Decamps, Anthony Herrel, Alexis Chaine, Violaine Llaurens, Vincent Debat

## Abstract

Predation is a powerful selective force shaping many behavioural and morphological traits in prey species. The deflection of predator attacks from vital parts of the prey usually involves the coordinated evolution of prey body shape and colour. Here, we test the deflection effect of hindwing tails in the swallowtail butterfly *Iphiclides podalirius*. In this species, hindwings display long tails associated with a conspicuous colour pattern. By surveying the wings within a wild population of *I. podalirius*, we observed that wing damage was much more frequent on the tails. We then used a standardised behavioural assay employing dummy butterflies with real *I. podalirius* wings to study the location of attacks by great tits *Parus major*. Wing tails and conspicuous coloration of the hindwings were struck more often than the rest of the body by birds. Finally, we characterised the mechanical properties of fresh wings and found that the tail vein was more fragile than the others, suggesting facilitated escape ability of butterflies attacked at this location. Our results clearly support the deflective effect of hindwing tails and suggest that predation is an important selective driver of the evolution of wing tails and colour pattern in butterflies.

## Introduction

Predation often affects the evolution of multiple morphological and behavioural traits in prey species. While many traits limiting predator attacks evolve, traits increasing survival after an attack have also been repeatedly promoted by natural selection [1]. Traits enhancing attack deflection, by attracting strikes towards a conspicuous body part, indeed limit damage to vital parts and increase escape probability [2]. The conspicuous coloration on the tails of some lizard species has been suggested to promote attacks on the tails, therefore limiting wounds on other parts of the body ([3],[4]). The attraction towards conspicuous tails can also be reinforced by striped body coloration, directing the attention of predators towards the tail [5]. In salamanders, defensive posture increases tail conspicuousness [6], suggesting that both body shape and colour, as well as behaviour, may contribute to the deflecting effect. The emergence of a deflecting effect may thus result from a joint evolution of several morphological and behavioural traits (reviewed in [7] for lizards). In butterflies, the joint evolution of hindwing tails and specific behaviour enhancing attack deflection has been shown in *Lycaenidae*. In these butterflies, the hindwings frequently display tiny tails, conspicuous colour patterns and a specific behaviour involving tails movements, hypothesised to mimic a head with moving antennae (the “false head effect”, [8],[9]). The ‘false-head’ tails of *Lycaenidae* are likely to deflect attacks away from vital parts [8]. Laboratory experiments with spiders indeed showed that *Calycopis cecrops* butterflies, displaying false-head hindwings, escaped more frequently than butterflies from other species where hindwings do not display such false-heads [10]. In Museum collections, the prevalence of individuals with symmetrically damaged hindwings, interpreted as beak marks of failed predator attacks, has been shown to be higher in *Lycaenidae* species with wing tails, as compared to species without a tail or with a less conspicuous colour pattern [11]. This suggests that the deflecting effect associated with hindwing tails might rely on the joint evolution of wing shape, colour pattern, and behaviour, promoted by the attack behaviour of predators relying on visual cues.

Such a deflecting effect may lead to the loss of the attacked body part, but with limited effect on survival. In lizards and salamanders, tails can be detached without severely impacting survival of the attacked animal (*i*.*e*. autotomy [12]; [13]). In butterflies, wing margins displaying eyespots are preferentially attacked (e.g. in *Bicyclus anynana* [14], in *Lopinga achine* [15]). The loss of wing margins and especially hindwing margins has a low impact on butterflies flying abilities [16] and may therefore have a limited impact on survival. Butterflies are indeed commonly observed flying in the wild with such wing damage [17]. The escape from predators after an attack might also be facilitated by enhanced fragility of the attacked parts of the wings. In *Pierella* butterflies for instance, Hill and Vaca (2004) [18] showed that the conspicuous areas of the hindwings are associated with increased fragility, which may facilitate the escape after a predation attempt directed at this specific wing area. Similarly, in small passerine bird species, the feathers located in the zone most prone to the predator attacks are easier to remove [19]. The evolution of specific body parts with increased fragility might thus be promoted by predation pressure, because they enhance prey survival after an attack.

The repeated evolution of hindwing tails in *Lepidoptera* could result from the selection exerted by predators on the evolution of traits that enhance deflection. The long, twisted wing tails of some *Saturniidae* moths have indeed been shown to divert bats from attacking moth bodies [20]. During flapping flight, the spinning tails indeed confuse the echolocation signal perceived by predators, thus diminishing strike efficiency [21]. The evolution of wing tails in moths is thus likely to be promoted by the sensory system of their nocturnal predators. The deflecting effect of wing tails has also been suggested in day-flying butterflies facing diurnal predators relying on visual cues, but has been tested only in the very specific case of the false-head wing tail of *Lycaenidae*.

A large number of other butterfly species with diurnal activities nevertheless display hindwing tails. Swallowtail butterflies are particularly well-known for their conspicuous, highly diversified hindwing tails [22], but the selection exerted by predators on the repeated evolution of these tails has never been formally investigated. Here, we tested whether the evolution of tails might be promoted by attack deflection, using the swallowtail species *Iphiclides podalirius* (Linné, 1758 *Lepidoptera, Papilionidae*) as a case-study. *I. podalirius* is a large palearctic butterfly with hindwings displaying long tails associated with a salient colour pattern: an orange eyespot and four blue lunules with strong UV reflectance [23]. The combination of hindwing tail and colour pattern is therefore very conspicuous (Figure 3), and especially for predators sensitive to UV reflection, such as songbirds [24]. Moreover, the four wings exhibit convergent black stripes over a pale background, contiguous between forewings and hindwings in resting position, pointing towards the anal edge. This may enhance the attraction of a predator to the posterior part of the hindwing [8]. To test whether the evolution of wing tails in this species may stem from selection promoting traits enhancing attack deflection, we performed a series of three complementary experiments.

First, we characterised the amount and location of damage on the wings of wild butterflies to test whether tails are more frequently lacking in surviving butterflies, possibly indicative of failed predation attempts. Second, we conducted experimental behavioural assays in captivity using an avian generalist predator, the Great tit *Parus major*, and dummy butterflies made with real *I. podalirius* wings, in order to investigate the location of attacks. We specifically tested whether attacks are more frequently directed towards the hindwing tails and associated colour pattern as compared to the rest of the butterfly body. Finally, we used a specific experimental set up to estimate the force needed to tear wings at different locations. Preferentially-attacked body parts are predicted to be more easily detached, as it would enhance the probability of escape of the butterfly after an attack [18]. This combination of experiments in controlled and under natural conditions provides a test for the role of predator deflection in the adaptive evolution of wing shape, wing colour pattern and wing resistance in swallowtail butterflies.

**Table 1:**
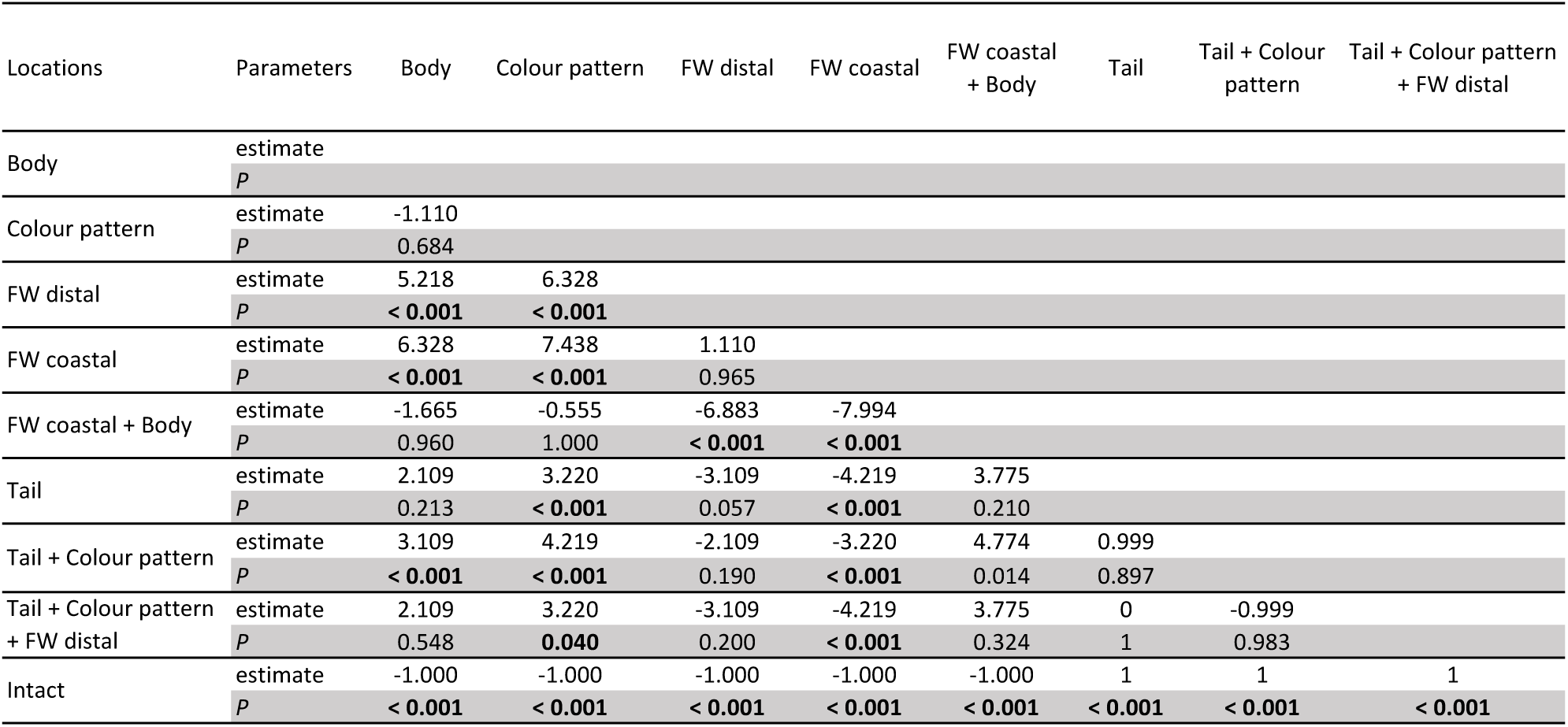
Post Hoc comparisons of bird strike numbers on the dummy butterflies between attack locations. Eight categories of location were defined: Body, FW coastal, FW distal, Colour pattern, Tail, FW coastal + Body, Tail + Colour pattern, Tail + Colour pattern + FW distal.

## Materials and Methods

### Field sampling

Field sampling of *I. podalirius* was performed in Ariege (France) during the summer of 2020 (Collection sites: 43°04′17.86″N, 01° 21′58.88″E; ca. 400 m a.s.l., and 43°03′50.94″N, 01° 20′40.95″E; ca. 400 m a.s.l.). We sampled a total of 138 wild individuals, with a large majority of males (132 males/6 females), likely reflecting the patrolling behaviour displayed by males (hill topping). After their capture, butterflies were euthanized by hypothermia and their wings stretched out and dried.

### Assessing the distribution of wing damage in the wild

The dorsal side of the forewings (FWs) and hindwings (HWs) of the field-sampled individuals was photographed in controlled LED light conditions (Nikon D90, Camera lens: AF-S Micro Nikkor 60 mm 1:2.8G ED). Out of the 138 wild butterflies collected, 65 exhibited wing damage. We studied the location of missing wing areas, distinguishing damage occurring on HW and FW, and reported the asymmetry of different types of damage (left and right damage with visually similar areas and positions were considered symmetric). A Pearson’s Chi-squared test with Yates’ continuity correction was used to test whether (1) damage was more often observed on hindwings than on forewings, and (2) damage on hindwings was more often asymmetric than damage on forewings.

To finely quantify the distribution of missing wing areas, we then digitised the wing outlines of the 65 damaged butterflies and 10 intact individuals as references. We defined 300 semi-landmarks equally spaced along the outline of both the left- and right-reflected FWs and HWs, using TpsDig2 [25]. The average shape of intact butterflies was obtained with TpsRelw, [25]), using a geometric morphometric approach ([26]; [27]). The wing outline of each damaged individual was then manually superimposed on the average shape of intact butterflies, in order to characterise the missing area of each damaged wing. A heat map was then obtained by summing up the occurrences of missing areas at each pixel throughout the sample of damaged individuals, using EBImage (R package; [28]), following [16]. The heat map was then plotted with autoimage (R package; [29]).

### Behavioural experiment with birds

We conducted an experiment to determine the location of attacks by birds on *I. podalirius* wings between October 2020 and January 2021 at the Station d’Ecologie Théorique et Expérimentale du CNRS, France (near the collection sites). Great tits were caught in mist-nets in the vicinity of the research station, ringed, and housed in individual indoor/outdoor cages (5m x 1m x 3m) and fed *ad libitum* with mealworms and sunflower seeds. After 2 days of habituation to captivity, we conducted behavioural experiments on two consecutive days during the three hours after sunrise while birds did not have access to sources of food other than dummy butterflies. The whole experiment was repeated three times using new birds for a total of 72 different birds tested. Capture of wild birds was performed under permits from the French ringing office (CRBPO, permit 13619 to A. Chaine). Capture and holding of birds from the wild was approved by the Région Midi-Pyrenées (DIREN, n°2019-s-09) in the Moulis experimental aviaries (Préfecture de l’Ariège, institutional permit n°SA-12-MC-054; Préfecture de l’Ariège, Certificat de Capacite, n°09-321 to A. Chaine).

We built 95 dummy butterflies, using actual wings of *I. podalirius* butterflies collected in the wild, glued on an artificial black cardboard body. The position of the glued wings corresponded to the natural position of butterflies at rest (Figure 1). A dummy was placed in each bird cage, about 1.5 m off the ground, using a wire fixed to the cage wall. This setting thus allowed the dummy to gently “flutter” in the middle of the cage, far enough from any perching site, to prevent close inspection by resting birds. The birds thus had to approach and potentially strike dummy butterflies while flying. Each cage was equipped with a camera filming continuously (Figure 1, S2). Two observers also monitored the 24 experimental cages: damaged dummies were replaced as soon as noticed by the observers, to maximise the number of attacks on intact butterflies. After each experimental session, the birds were fed *ad libitum* until nightfall to minimise the stress generated by the experiment. The whole experiment was repeated for 2 consecutive days.

**Figure 1:**
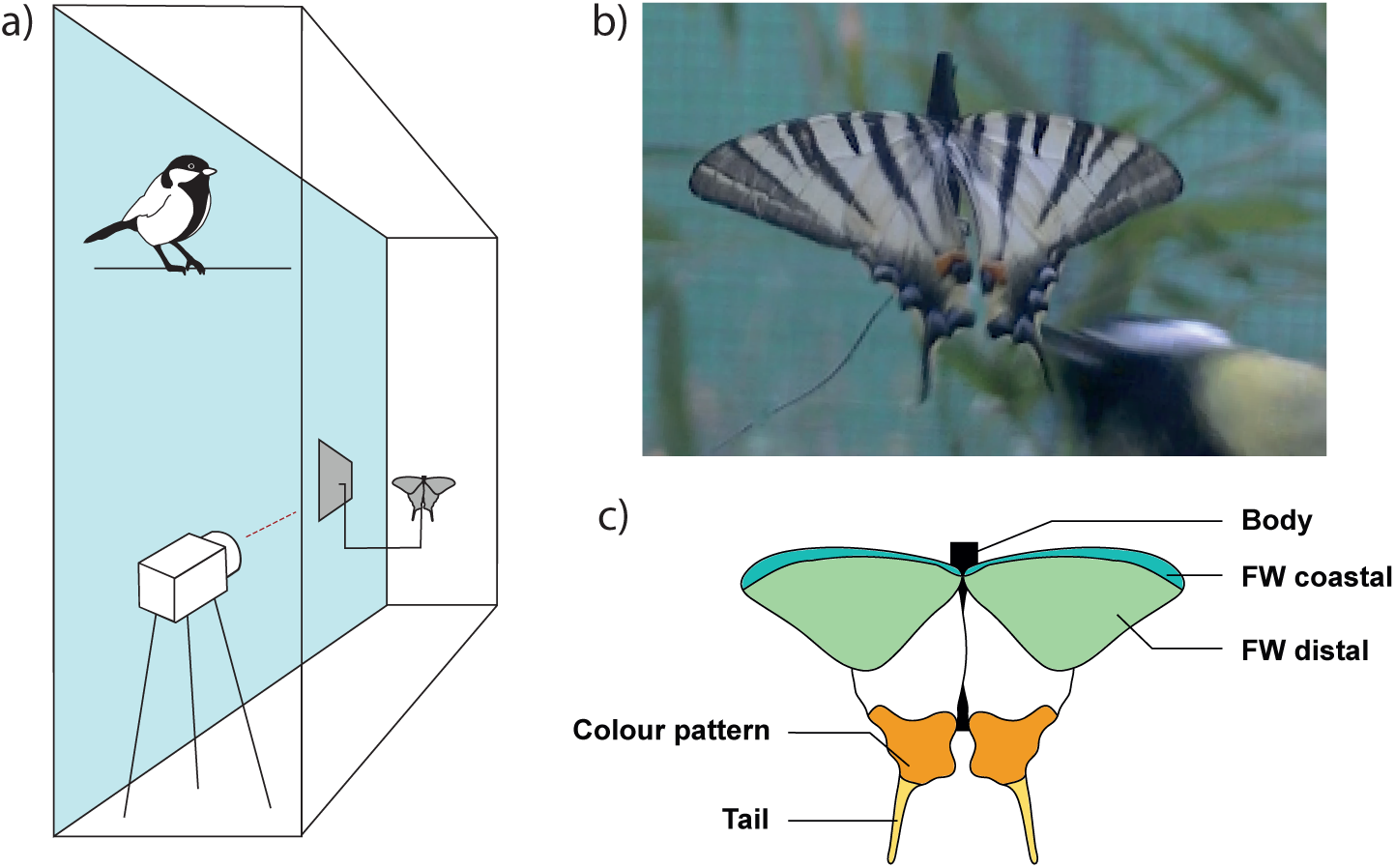
Experimental setup for behavioural assay with wild-caught great tits. (a) Each experimental cage (5m x 1m x 3m) was equipped with a video camera filming continuously. A butterfly dummy was fixed to the wall at about 1.5 m off the ground using a wire far enough from any perching site to prevent close inspection by the birds not in flight. (b) Picture of a butterfly dummy struck by a bird. (c) Schematic of a dummy butterfly composed of four real wings glued on an artificial black cardboard body. Five locations could be targeted by birds: Body, FW coastal, FW distal, Colour pattern, and Tail. Photograph of the setup in Supplementary materials S2.

Analyses of videos recorded during the experiments were used to count the exact number of strikes performed by each bird on each dummy butterfly. Each strike was defined as a single touch of the beak on the dummy butterfly. The films were also used to assess the precise location of each strike on the butterfly body. Five categories of strike location were defined: body, coastal part of the forewing, distal part of the forewing, hindwing colour pattern and hindwing tail (see Figure 1). In some cases, the strike affected several locations at once. These “combined” locations were considered as separate categories, leading to a total of 8 possible targeted locations (see Figure 3). Because a dummy could be attacked several times before it was replaced, we also recorded the order of each strike performed by the tested bird on the given dummy.

We first tested whether strikes occurred more often on the hindwings than on the forewings, using a Pearson Chi-squared test with Yates’ continuity correction. To test whether the different parts of the wings were equally prone to attack, we applied a generalised binomial regression model for the probability of attack, using strike location and strike rank order as effects and considering all specimens and sessions (including birds that did not attack). An analysis of variance was then applied (ANOVA type II, function “Anova’’ package “car “, [30]). To perform pairwise comparisons on the location categories, we finally performed a series of post hoc tests (function “tukey_hsd” package “rstatix “, [31]).

### Mechanical resistance of the wings

#### Experimental sample

We tested mechanical resistance of the different wing parts on 28 fresh *I. podalirius* butterflies (21 females and 7 males) obtained as pupae from a commercial supplier (Worldwide Butterflies Ltd). After emergence, individuals were placed in individual cages to allow proper unfolding and drying of the wings, then placed in entomological envelopes to avoid wing damage. The butterflies were fed once a day with a mixture of water and honey, and maintained for 11 to 20 days depending on the time between emergence and the start of the experiments. Experiments were performed on freshly-killed individuals to limit the effect of wing drying on mechanical properties [32]. In order to test whether the tails are more fragile, we compared the mechanical resistance of different regions of the wings (Figure 2). Specifically, we contrasted the vein located within the hindwing tail (M3H vein), with another hindwing vein located outside the colour pattern area (R5H vein). We also included the two developmentally-homologous veins on the forewing (M3F and R5F; [33]). For each butterfly, the experiment was conducted on one hindwing and one forewing. The four veins were measured in a randomised order to avoid any bias caused by the deformation of the wings due to previous tearing.

**Figure 2:**
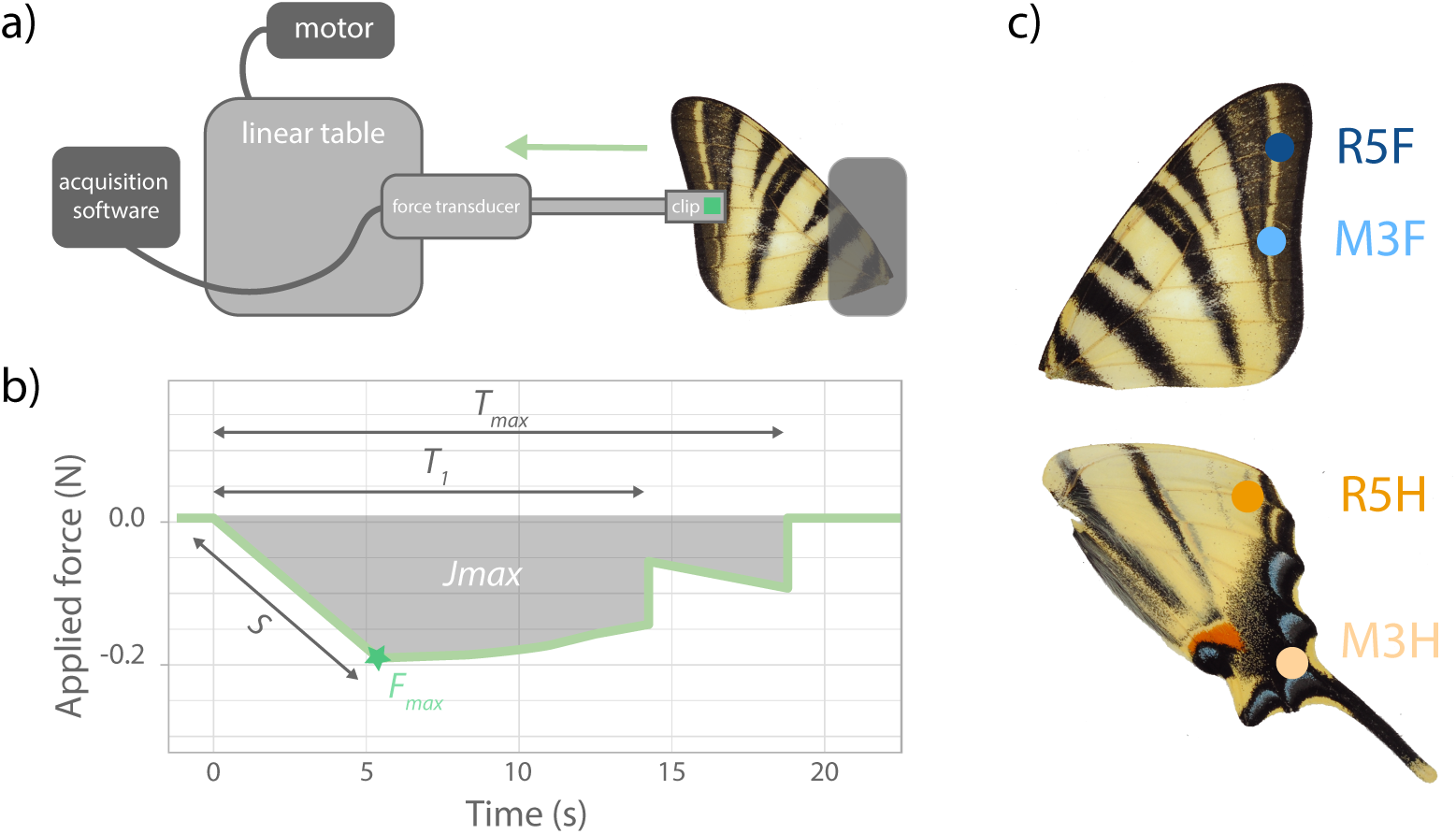
Experimental setup designed to estimate the strength needed to break wings at different locations. (a) A custom setup was built, composed of a mobile part (clip + force transducer + linear table) exerting traction on the wing and a fixed part, holding the wing. (b) Summary variables derived from force profile: *F*_*max*_ (the maximum force exerted on the vein), *T*_*1*_ (the time to the first break), *T*_*max*_ (the time to the complete rupture of the vein), *S* the slope of the curve (estimating the stiffness of the wing) and the area under the curve *J*_*max*_ (indicating the impulse). (c) Locations of the four measured points with attachment on a vein at 3mm from the edge of the wings. Hindwing tail resistance is measured at the point M3H. Photograph of the setup in Supplementary materials S3.

#### Experimental setup

As wing parts involved in predator deflection are expected to be particularly fragile, we designed a custom experimental set-up adapted from Hill and Vaca (2004) [18] and De Vries (2002) [34] to specifically estimate the mechanical resistance of different parts of the wings. When a bird catches the wing of a butterfly, the force exerted by the beak and the opposed escape movement of the butterfly likely induce tensile stresses on the wing. We thus compared the mechanical response of the different wing veins to a tensile force exerted in the direction of the vein, away from the body (see Figure 2). Our set up was composed of a fixed part holding the wing and of a mobile part exerting traction on the wing (Figure 2, S3). This mobile part was connected to the wing using a flattened and filed alligator clip with a squared 9 mm^2^ piece of rubber ensuring a soft and standardised contact with the wing. For each measurement, the clip was fixed at 3mm from the edge of the wing. This clip was then connected to a piezo-electric force transducer (Kistler 9217A type 9207 Serian number: 1275844), connected to a charge amplifier (Kistler type 5011). The force was fixed on a linear table controlled by a motor (RS PRO, 12V dc, 2400 gcm), allowing constant traction. The charge from the force transducer was measured by the amplifier and sent to a Biopac AD unit. Forces were captured and analysed using AcqKnowledge software (version 4.1, BIOPAC Systems, Inc.).

The variation of the force through time, from the onset of the motor to the total rupture of the wing was recorded for each trial. These response curves were first smoothed using a lowpass filter set at 20HZ. Five summary variables were extracted from the response curve (Figure 2): (1) the maximum force exerted on the vein (estimating the maximum strength of the vein, noted *F*_*max*_), (2) the time to the first break (*T*_*1*_; shown by the first abrupt decrease in force), (3) the time to the complete rupture of the vein (*T*_*max*_; when the force returns to zero), (4) the slope (*S*) of the curve between the beginning of the pull and the point of maximum force (estimating the stiffness of the wing - see Supplementary materials S1) and (5) the impulse required for the complete rupture of the vein (*J*_*max*_), assessed by the area under the curve. The forces were measured in Newtons (N) - note the force takes negative values since we measured a tensile force.

#### Statistical analysis

The five mechanical parameters measured on the different veins were then compared using linear mixed models using wing (forewing *vs*. hindwing) and vein (M3 *vs*. R5) as fixed effects, while butterfly ID, sex and the date of measurement session were set as random variables (function “lmer”, package “lmerTest”, [35]). The date of measurement session was added to account for potential differences in heat and humidity across session possibly affecting the wing mechanical properties. For *J*_*max*_, there was some evidence that the wing and vein effects interacted. We thus modified the model to directly account for the four modalities of the vein effect (R5F, M3F, R5H, M3H). We analysed all models with a type III analysis of variance. All statistical analyses were carried out in R version 4.0.3 (R Core Team 2020).

## Results

### Wing damage mostly affects the tails

We hypothesised that a deflection effect should result in a higher proportion of wing damage on the deflecting wing areas in the wild. To test this hypothesis, we studied the location of wing damage in a natural population of *I. podalirius*. Among all wild individuals collected, 47.1% had wing damage. Forewings were less often damaged than hindwings (22.31% and 85.38% respectively; *χ*^*2*^ = 101.54, *df* = 1, *P* < 0.001). The frequency of individuals with missing hindwing tails in the wild was especially high: all 65 damaged individuals had at least one tail damaged (out of 130 wings tested, 82.3% had tail damaged). This result is illustrated by the heatmap (Figure 5). Furthermore, damage on the hindwings were more often asymmetrical (78.46%) than damage on the forewings (24.62%) (*χ*^*2*^ = 35.603, *df* = 1, *P* < 0.001).

**Figure 3:**
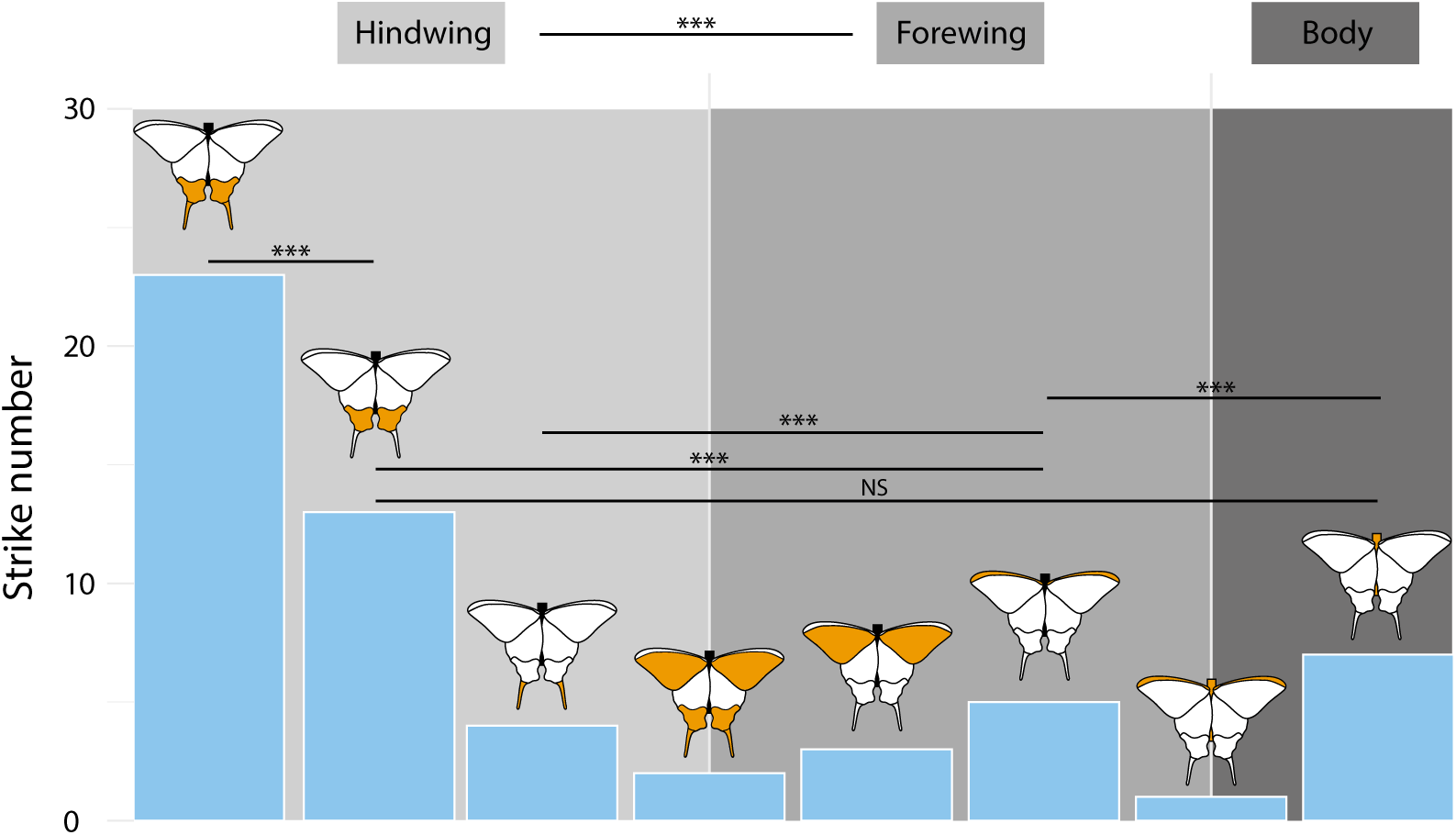
**Locations of bird strikes on the dummy butterflies**, recorded during six experimental sessions on 72 captured *Parus major* using butterfly dummies built with real wings of *I. podalirius*. A total of 59 strikes were recorded. Each category is defined by the location targeted by a bird in a single strike and represented in orange on each associated butterfly scheme. Only essential statistical comparisons are represented; see details in Table 1. Video in Supplementary movies 1-2.

**Figure 4:**
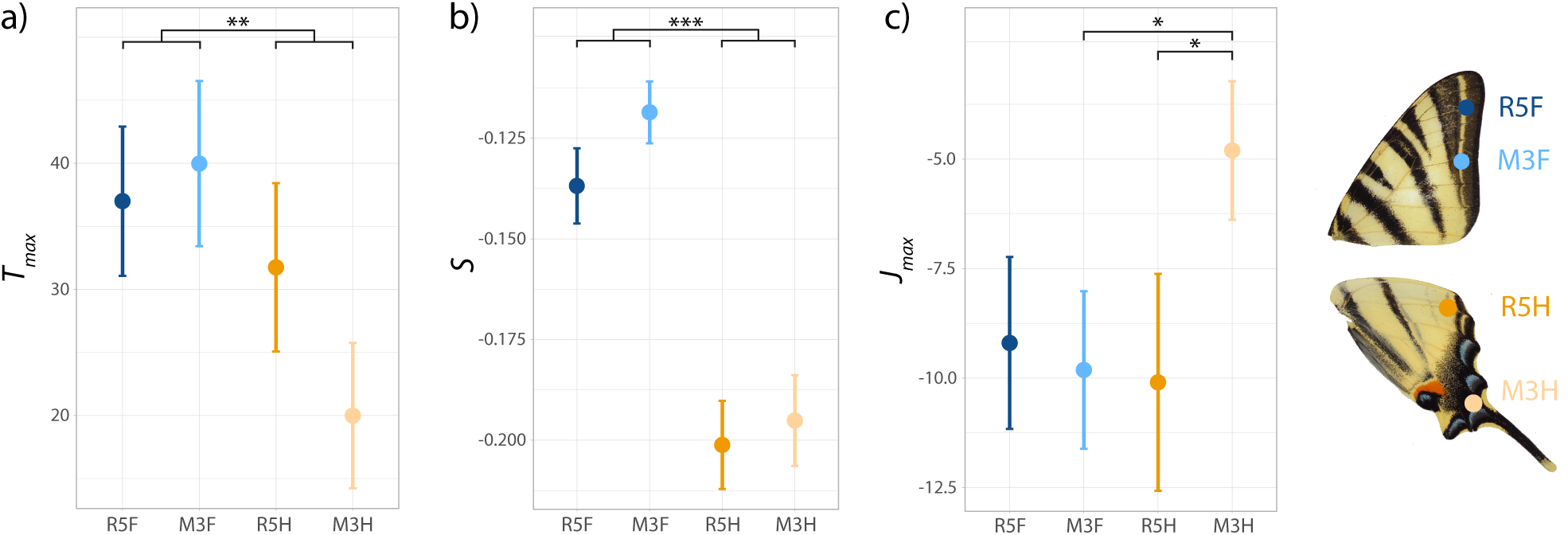
Variation in mechanical resistance in different areas of the forewing and hindwings of fresh *I podalirius* samples (n=28). On each of the 28 butterflies, four locations were studied, corresponding to four different veins (R5F, M3F, R5H, M3H). Means and standard errors are indicated as well as significant differences between locations. Three mechanical variables per wing location are reported (a) *T*_*max*_ (the time to the complete rupture of the vein). (b) *S*, the slope of the curve (estimating the stiffness of the wing) and (c) *J*_*max*_, the area under the curve (a measure of impulse).

**Figure 5:**
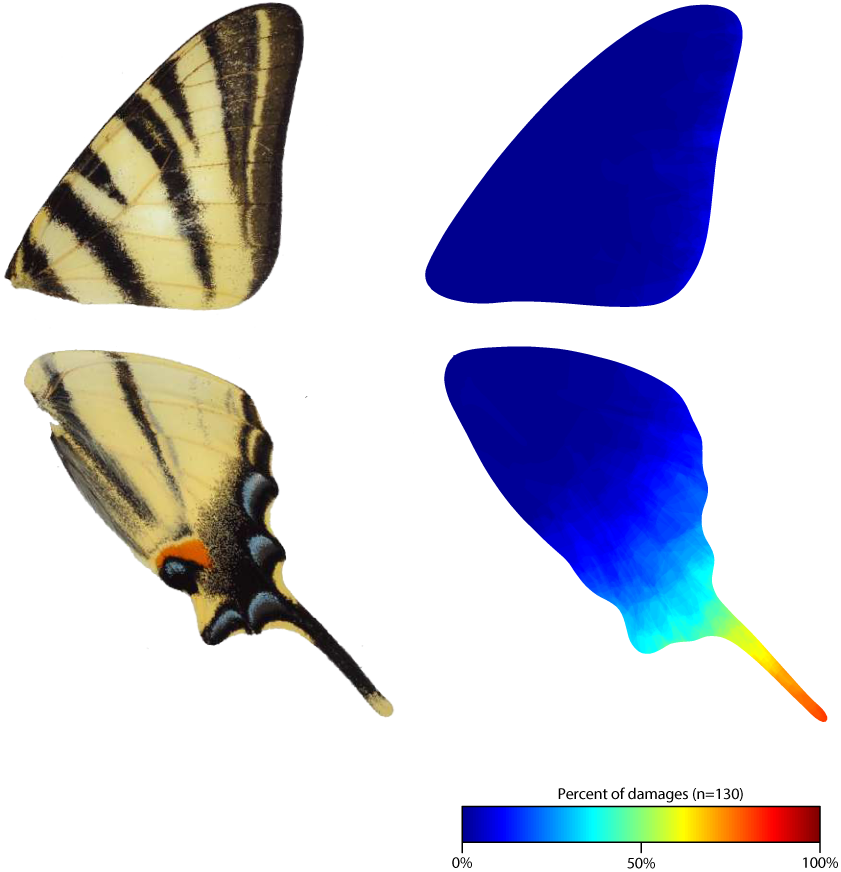
Heatmap describing the spatial distribution of wing damage on a sample of wild *I. podalirius*. Left: Photograph of *I. podalirius* wings. Right: Proportion of naturally damaged wing locations. Data for left and right wings were pooled for each pair of wings (65 individuals, so 130 forewings and 130 hindwings). The most frequently damaged areas are shown in red, while intact areas are shown in blue (see colour scale).

### Behavioural experiments with birds reveal preferential attacks on hindwing tail and colour pattern

Using the behavioural assays carried out with great tits, we investigated whether the attacks on dummy butterflies were directed towards the posterior part of the hindwings (Figure 3), as expected under the hypothesis of a deflecting effect induced by the butterfly morphology. Among the 72 birds tested, only 17 attacked the dummy butterflies, resulting in 65 recorded strikes. Because some strikes occurred outside of the field of view of the camera, the targeted part of the dummy could be determined in only 59 of these strikes. The hindwings were more often targeted by the birds (43 strikes; 72.9%) than the forewings (16 strikes; 27.1%) (*χ*^*2*^ = 12.36, *df* = 1, *P* < 0.001). The probability of attack strongly depended on the wing location (*LR χ*^*2*^ = 141.21, *df* = 8, *p* < 0.001): there was strong evidence that strikes jointly targeting the tail and the colour pattern of the hindwings (23 attacks; 39%) were more frequent than strikes on any other body part (see detailed statistical tests in Table 1). In contrast, no evidence for an effect of the attack ranking on attack probability was found (see detailed statistical tests in Supplementary table 1).

### Hindwings and in particular hindwing tails are more easily damaged

We then tested whether the hindwing region with the tail and conspicuous colour patterns is more fragile than the rest of both wings, as expected if they are involved in a deflecting effect. There was a strong evidence that time to first rupture (*T*_*1*_) and the time to total rupture (*T*_*max*_) were lower in hindwings veins than forewing veins (Figure 4, see statistical tests in Table 2). *J*_*max*_, the impulse required to fully rupture the vein (as assessed by the area under the response curve; Figure 2) was smaller for the hindwing tail vein (M3H) than for any other veins (M5H: *t* = -2.42; *P* = 0.019; M3F: *t* = -2.48; *P* = 0.016; M5F: *t* = -1.88, *P* = 0.06). The slope of the force profile, *S*, reflects the stiffness of the wing: the greater the slope, the stiffer the veins (equations in Supplementary materials S1). There was strong evidence that hindwing veins had higher force profile slopes than forewing veins (Figure 4, details in Table 2), indicating that they are stiffer. Finally, a weak evidence for a lower F_max_ (maximum force applied to the vein) in the hindwings than in the forewings was found (*F* = 3.11; *P* = 0.082, Table 2).

**Table 2:** Summary of the linear mixed-effects models describing the effect of Wing (forewing/hindwing) and Vein (M3/R5) on the five mechanical parameters measured during the mechanical resistance experiment in different areas of the forewing and hindwings of fresh *I podalirius* samples (n=28): *F*_*max*_ (the maximum force exerted on the vein), *T*_*1*_ (the time to the first break), *T*_*max*_ (the time to the complete rupture of the vein), *S* (estimating the stiffness of the wing), *J*_*max*_ (the impulse required for the complete rupture of the vein). These five models were analysed with a type III analysis of variance.

|  | Wing |  |  | Vein |  |  | Wing:Vein |  |  |
| --- | --- | --- | --- | --- | --- | --- | --- | --- | --- |
|  | *df* | *F-value* | *p-value* | *df* | *F-value* | *p-value* | *df* | *F-value* | *p-value* |
| $F_{max}$ | 75.8500 | 3.1006 | 0.0823 | 75.6680 | 0.0180 | 0.8937 | 75.791 | 1.3071 | 0.2565 |
| $T_1$ | 44.4160 | 17.7794 | < **0.001** *** | 44.1520 | 0.8624 | 0.3581 | 44.793 | 2.7123 | 0.1066 |
| $T_{max}$ | 67.7270 | 7.9865 | **0.0062** ** | 64.3680 | 1.1973 | 0.2779 | 63.458 | 2.1050 | 0.1517 |
| $S$ | 69.0870 | 60.4044 | < **0.001** *** | 68.9320 | 1.3691 | 0.2460 | 70.49 | 0.3050 | 0.5825 |
| $E_{max}$ | 64.036 | 1.8635 | 0.1770 | 58.537 | 2.0436 | 0.1582 | 57.844 | 4.1549 | **0.04609** * |

## Discussion

Our multi-pronged approach combining behavioural experiments, biomechanical measurements and survey in natural population provide strong evidence of a deflecting effect of hindwing tails in *I. podalirius*, opening new research avenues on the predation pressures involved in the evolution of tails in butterflies.

### Adaptive evolution of hindwing tails promoted by predator behaviour

Our behavioural trials showed that attacks by great tits on *I. podalirius* are highly biased towards the hindwing tails and colour pattern. This provides strong support for a deflective effect generated by both colour pattern and tail on predators. Interestingly, only a small fraction of the tested birds actually attacked the dummies. This could suggest that *I. podalirius* butterflies are not the usual prey consumed by great tits [36], especially during the season when the tests were carried out (late fall and winter), where they mostly rely on seeds rather than on insects. Our behavioural experiments are thus relevant for the behaviour of generalist predators that are probably naïve to the phenotypes of the tested butterflies, a likely situation in nature, as no specialist predator is known for *I. podalirius*. Some of the birds nevertheless repeatedly attacked the posterior area of the hindwings, consecutively targeting the two tails, showing a particularly strong interest for this location (see Supplementary movie 1).

Birds typically flew above the butterflies, patrolling the cage at 3m from the ground and dummies had their tails oriented towards the ground about 1.5m from the ground. The high frequency of attacks on the tails therefore did not result from an easier access to the tails due to a positional bias. To the contrary, birds adjusted their trajectory to attack from below (see Supplementary materials S4; Supplementary movie 2), suggesting they were specifically targeting the tails. The combination of tails and associated conspicuous colour patterns is thus probably very attractive to predators, inducing the observed pattern of attack locations. Given the tested birds preferentially attacked the distal area of the hindwing, we would expect that this wing area should be easier to tear off. Such enhanced fragility would facilitate the butterfly escape and may thus be promoted by natural selection generated by the behaviour of birds.

Our analysis of the mechanical resistance of wing veins indeed shows that hindwing veins, and especially the vein located within the tail, are less resistant to the application of a tensile force and break sooner than forewing veins. Whether the measured difference in strength would have a significant impact during a predator attack is unknown but the forces tested are relevant to the type of strikes observed in our behavioural experiment. The enhanced fragility of the hindwing vein located within the tail is thus consistent with the deflection hypothesis. It should increase escape probability, while preserving the integrity of the wing and reducing aerodynamic costs. Interestingly, in *Pierella* butterflies, the conspicuous white patch of the hindwing found to have an increased fragility by Hill & Vaca (2004) [18] contains the M3H vein, *i*.*e*. the vein located within the tail in *I. podalirius*, that was found to be the stiffest and to break the earliest in our study. The M3H vein could have an enhanced fragility in many butterfly species, therefore promoting the evolution of conspicuousness in these wing areas, enhancing survival after an attack. The evolution of such association should especially be favoured if butterflies missing such wing area still survive in the wild.

The large abundance of tailless *I. podalirius* flying in the wild indeed testifies to the limited aerodynamic consequences of such damage. Tail loss does not prevent these damaged butterflies from performing their typical hill-topping behaviour and is thus likely to have a limited impact on their fitness. The distribution of damage across the wings in the natural population of *I. podalirius* also confirms that hindwing tails are more prone to attack than any other part of the wing (Figure 5). Inferring predation from butterfly wing damage alone can be misleading because damage can stem from a diversity of sources, including interactions with conspecifics ([37]; [38]) or collision with obstacles ([39]; [16]). However, the pattern we found is still consistent with an increased attack rate on hindwing tails. While damage due to collisions should be symmetrical as seen on forewings, the prevalence of asymmetric damage on the tails of *I. podalirius* matches the hypothesis of predator attacks during flight or when butterflies are at rest, typically perching on high branches with their wings wide open (Figure 1). This also suggests that symmetry in the tail is not critical for aerodynamics. Our survey in natural population thus reinforces the evidence for the adaptive evolution of tail and colour pattern in *I. podalirius*, where the benefits in terms of escape ability may exceed the costs of wing damage.

Considered together, (1) the strong prevalence of the attacks on the hindwing tails and associated colour patterns, (2) the reduced strength or the corresponding parts of the wings and (3) the very high incidence of natural wing damage on the tails, provide evidence for the adaptive evolution of hindwings tails in *I. podalirius via* a deflecting effect of predator attacks. The effect of attack deflection on the evolution of wing tails in day-flying butterflies has only been demonstrated in the peculiar case of false head morphology in *Lycaenidae* ([8]; [9]). Our study suggests that predation can be a major selective pressure involved in the evolution of hindwing tails in butterflies. Hindwing tails have evolved multiple independent times throughout the diversification of butterflies and are associated with an important diversity of colour patterns ([40]). Our results thus open the question of the evolution of different traits involved in predator deflection, namely hindwing shape, fragility and colour patterns, as well as behaviour, jointly forming an adaptive syndrome.

### Adaptive syndrome of predation deflection

In our experiments with birds, tails alone were targeted in a large proportion of the trials, but most attacks involved a combination of the tails and associated colour pattern. This strongly suggests that the visual effect triggering attack deflection in *I. podalirius* is jointly induced by the tails and the colour pattern, including the blue marks and the orange eyespots on the hindwing, and possibly the black stripes pointing at the tails. The deflection effect therefore probably relies on the evolution of a series of traits, including wing shape, wing colour pattern and wing mechanical resistance. The joint *vs*. sequential nature of the evolution of these different traits is largely unknown and might depend on the developmental and genetic bases of the traits involved in deflective syndromes, as well as the different selection pressures acting on each of those traits.

Associations between hindwing tails and peculiar colour patterns promoted by predation pressure have been described for butterfly species involved in Batesian mimicry. In *Papilio memnon*, for instance, some females display hindwing tails and red coloration resembling the toxic species *Pachliopta coon* on the Malay peninsula while other females have no tail and an alternative yellow colour pattern mimicking *Troides helena* in Northern Borneo [41]. These two traits are controlled by different loci and the linkage disequilibrium between these loci might have been promoted by the selective advantages brought by mimicry [42]. Nevertheless, the association between well-developed tails and conspicuous colour elements is not universal in Papilionidae: for example, *Papilio ulysses* tails and surrounding wing parts are completely black, while in *Papilio demodocus*, conspicuous distal eyespots are observed in tailless hindwings. Shared developmental pathways in wing shape and colour pattern might promote their joint evolution, so that the emergence of deflective syndromes can be facilitated in some lineage. Alternatively, species ecology might trigger strong selection promoting linkage disequilibrium between loci controlling traits enhancing deflection.

The combined evolution of traits limiting predation also frequently extends to behaviour. Whether the behaviour emerges before or after the evolution of morphological traits involved in deflection is an open question. In *I. podalirius*, the perching position with wings wide open possibly enhances the deflecting effect provided by hindwing tails but might have been promoted for its effect on thermoregulation [43] before the evolution of tails. Adaptive syndromes involving the evolution of both morphological and behavioural traits promoted by predator behaviour have been observed in other *Lepidoptera*. In some species, hidden conspicuous coloration can be suddenly uncovered when threatened by a predator, inducing a startling effect (*e*.*g. Catocala nupta*, [44]). The evolutionary sequence of these behavioural and morphological traits has been investigated experimentally by testing the deterring effect of both traits independently. These experiments suggest that behavioural changes might have preceded the evolution of conspicuous coloration, because sudden movements can be sufficient to induce strong deterrence ([45]). Whether a similar ‘behaviour first’ evolutionary sequence is involved in the evolution of deflective syndromes should be investigated.

Important selective trade-offs between predator deflection and flight abilities might also influence the evolution of deflective syndromes in *Lepidoptera*, therefore constraining wing areas involved in such syndromes. Anteromotorism being a shared characteristic of butterflies [46], hindwing fragility might be ancestral, and conspicuous marks might have secondarily been favoured on these weaker wings. In Papilionidae, hindwing shape is indeed strikingly more diversified than forewing shape [22] in agreement with lower aerodynamic constraints on the hindwings. The study of aerodynamic forces applied to an artificial model of a butterfly with tails suggests that hindwing tails increase the lift of the butterfly during gliding [47]. Preservation of flight capacity through the maintenance of tail integrity, and in particular a sufficient strength to withstand the pressure forces applied during flapping, could act as an evolutionary trade-off with the selection of mechanical weakness. The selective pressures acting on each of the traits involved in these deflective syndromes should now be studied independently and compared in species with contrasted ecologies and levels of phylogenetic proximity to determine the evolutionary forces involved in the emergence of deflective syndromes.

## Conclusions

The diversity of wing tails observed in *Lepidoptera* suggests they have evolved multiple times, therefore raising the question of the selective pressures involved. Based on our combined analysis of natural wing damage, biomechanical resistance of the wings, and behavioural interactions with bird in the species *I. podalirius*, we provide evidence for an effect of natural selection exerted by predators on hindwing tail evolution, promoting traits enhancing attack deflection away from the vital body parts. Our study therefore opens up new research avenues on the relative effect of predation pressure *vs*. other selective forces involved in the evolution of hindwing tails in butterflies. We also highlight that such a deflective effect may have emerged from a sequential evolution of a suite of traits, including wing shape, wing colour patterns, and wing mechanical properties. These questions should stimulate new research on the developmental and selective origin of the traits involved in deflective syndromes in various butterfly species.

## Supporting information

Supplementary materials S1: Physical characterisation of the tensile strength of the wing (from [48]).

Supplementary materials S2: Photograph of the experimental setup for behavioural assay with wild-caught great tits.

Supplementary materials S3: Photograph of the experimental setup to estimate the force needed to tear the wings at different locations.

Supplementary materials S4: Strike trajectories of birds (n=59). We tested whether strikes occurred more often with a bottom-up trajectory than a top-

Supplementary table 1: Post Hoc comparisons of bird strike numbers on the dummy butterflies between rank order of attacks. Each strike was characteris

Supplementary movie 1: Video of three sequential strikes performed by a great tit on a dummy butterfly. Strikes are shown at normal speed then slowed

Supplementary movie 2: Video of one strike performed by a great tit on a dummy butterfly. Strike is shown at normal speed then slowed down 10 times.

## Acknowledgements

We would like to thank Camille Le Roy for his advice on the analysis of wing damage, Thomas Crouchet for his assistance during the behavioural assay and Ramiro Godoy-Diana and Roméo Antier for their advice on biomechanics and material deformation. Aviary work was supported by ANR-SoCo to A. Chaine and the Laboratoire d’Excellence (LABEX) entitled TULIP (ANR-10-LABX-41).

## Legends

**Supplementary table 1: Post Hoc comparisons of bird strike numbers on the dummy butterflies between rank order of attacks**. Each strike was characterised by its rank order, from 1 to 7.

**Supplementary materials S1: Physical characterisation** of the tensile strength of the wing (from [48]).

**Supplementary materials S2: Photograph of the experimental setup** for behavioural assay with wild-caught great tits.

**Supplementary materials S3: Photograph of the experimental setup** to estimate the force needed to tear the wings at different locations.

**Supplementary materials S4: Strike trajectories of birds (n=59)**. We tested whether strikes occurred more often with a bottom-up trajectory than a top-down trajectory using a Pearson Chi-squared test. Bottom-up trajectories (n=33) are more frequent than top-down (n=19) (*χ*^*2*^ = 6.3333, *df* = 1, *P* = 0.012).

**Supplementary movie 1: Video of three sequential strikes** performed by a great tit on a dummy butterfly. Strikes are shown at normal speed then slowed down 10 times.

**Supplementary movie 2: Video of one strike** performed by a great tit on a dummy butterfly. Strike is shown at normal speed then slowed down 10 times.

## Bibliography

1. Bateman AW, Vos M, Anholt BR. 2014 When to Defend: Antipredator Defenses and the Predation Sequence. Am. Nat. 183, 847–855. (doi:10.1086/675903)

2. Ruxton GD, Sherratt TN, Speed MP. 2004 Avoiding attack: the evolutionary ecology of crypsis, warning signals, and mimicry. Oxford; New York: Oxford University Press.

3. Watson CM, Roelke CE, Pasichnyk PN, Cox CL. 2012 The fitness consequences of the autotomous blue tail in lizards: an empirical test of predator response using clay models. Zoology 115, 339–344. (doi:10.1016/j.zool.2012.04.001)

4. Guidi R dos S, São-Pedro V de A, da Silva HR, Costa GC, Pessoa DMA. 2021 The trade-off between color and size in lizards’ conspicuous tails. Behav. Processes 192, 104496. (doi:10.1016/j.beproc.2021.104496)

5. Murali G, Kodandaramaiah U. 2016 Deceived by stripes: conspicuous patterning on vital anterior body parts can redirect predatory strikes to expendable posterior organs. R. Soc. Open Sci. 3, 160057. (doi:10.1098/rsos.160057)

6. Myette AL, Hossie TJ, Murray DL. 2019 Defensive posture in a terrestrial salamander deflects predatory strikes irrespective of body size. Behav. Ecol. 30, 1691–1699. (doi:10.1093/beheco/arz137)

7. Arnold EN. 1984 Evolutionary aspects of tail shedding in lizards and their relatives. J. Nat. Hist. 18, 127–169. (doi:10.1080/00222938400770131)

8. Robbins RK. 1981 The ‘False Head’ Hypothesis: Predation and Wing Pattern Variation of Lycaenid Butterflies. Am. Nat. 118, 770–775. (doi:10.1086/283868)

9. Wourms MK, Wasserman FE. 1985 Butterfly wing markings are more advantageous during handling than during the initial strike of an avian predator. Evolution 39, 845–851. (doi:10.1111/j.1558-5646.1985.tb00426.x)

10. Sourakov A. 2013 Two heads are better than one: false head allows Calycopis cecrops (Lycaenidae) to escape predation by a Jumping Spider, Phidippus pulcherrimus (Salticidae). J. Nat. Hist. 47, 1047–1054. (doi:10.1080/00222933.2012.759288)

11. Novelo Galicia E, Luis Martínez MA, Cordero C. 2019 False head complexity and evidence of predator attacks in male and female hairstreak butterflies (Lepidoptera: Theclinae: Eumaeini) from Mexico. PeerJ 7, e7143. (doi:10.7717/peerj.7143)

12. Beneski JT. 1989 Adaptive Significance of Tail Autotomy in the Salamander, Ensatina. J. Herpetol. 23, 322. (doi:10.2307/1564465)

13. Cooper WE. 1998 Reactive and anticipatory display to deflect predatory attack to an autotomous lizard tail. Can. J. Zool. 76, 4.

14. Chan IZW, Ngan ZC, Naing L, Lee Y, Gowri V, Monteiro A. 2021 Predation favours Bicyclus anynana butterflies with fewer forewing eyespots. Proc. R. Soc. B Biol. Sci. 288, 20202840. (doi:10.1098/rspb.2020.2840)

15. Olofsson M, Vallin A, Jakobsson S, Wiklund C. 2010 Marginal Eyespots on Butterfly Wings Deflect Bird Attacks Under Low Light Intensities with UV Wavelengths. PLoS ONE 5, e10798. (doi:10.1371/journal.pone.0010798)

16. Le Roy C, Cornette R, Llaurens V, Debat V. 2019 Effects of natural wing damage on flight performance in Morpho butterflies: what can it tell us about wing shape evolution? J. Exp. Biol. 222, jeb204057. (doi:10.1242/jeb.204057)

17. Molleman F et al. 2020 Quantifying the effects of species traits on predation risk in nature: A comparative study of butterfly wing damage. J. Anim. Ecol. 89, 716–729. (doi:10.1111/1365-2656.13139)

18. Hill RI, Vaca JF. 2004 Differential Wing Strength in Pierella Butterflies (Nymphalidae, Satyrinae) Supports the Deflection Hypothesis. Biotropica 36, 362–370. (doi:10.1111/j.1744-7429.2004.tb00328.x)

19. Møller AP, Nielsen JT, Erritzøe J. 2006 Losing the last feather: feather loss as an antipredator adaptation in birds. Behav. Ecol. 17, 1046–1056. (doi:10.1093/beheco/arl044)

20. Barber JR, Leavell BC, Keener AL, Breinholt JW, Chadwell BA, McClure CJW, Hill GM, Kawahara AY. 2015 Moth tails divert bat attack: Evolution of acoustic deflection. Proc. Natl. Acad. Sci. 112, 2812–2816. (doi:10.1073/pnas.1421926112)

21. Rubin JJ, Hamilton CA, McClure CJW, Chadwell BA, Kawahara AY, Barber JR. 2018 The evolution of anti-bat sensory illusions in moths. Sci. Adv., 10.

22. Owens HL, Lewis DS, Condamine FL, Kawahara AY, Guralnick RP. 2020 Comparative Phylogenetics of Papilio Butterfly Wing Shape and Size Demonstrates Independent Hindwing and Forewing Evolution. Syst. Biol. 69, 813–819. (doi:10.1093/sysbio/syaa029)

23. Gaunet A, Dincă V, Dapporto L, Montagud S, Vodă R, Schär S, Badiane A, Font E, Vila R. 2019 Two consecutive Wolbachia -mediated mitochondrial introgressions obscure taxonomy in Palearctic swallowtail butterflies (Lepidoptera, Papilionidae). Zool. Scr. 48, 507–519. (doi:10.1111/zsc.12355)

24. Cuthill IC, Partridge JC, Bennett ATD, Church SC, Hart NS, Hunt S. 2000 Ultraviolet Vision in Birds. In Advances in the Study of Behavior (eds PJB Slater, JS Rosenblatt, CT Snowdon, TJ Roper), pp. 159–214. Academic Press. (doi:10.1016/S0065-3454(08)60105-9)

25. Rohlf F. 2015 The tps series of software. Hystrix Ital. J. Mammal. 26. (doi:10.4404/hystrix-26.1-11264)

26. Adams DC, Rohlf FJ, Slice DE. 2004 Geometric morphometrics: Ten years of progress following the ‘revolution’. Ital. J. Zool. 71, 5–16. (doi:10.1080/11250000409356545)

27. Bookstein FL. 1997 Landmark methods for forms without landmarks: morphometrics of group differences in outline shape. Med. Image Anal. 1, 225–243. (doi:10.1016/S1361-8415(97)85012-8)

28. Pau G, Fuchs F, Sklyar O, Boutros M, Huber W. 2010 EBImage--an R package for image processing with applications to cellular phenotypes. Bioinformatics 26, 979–981. (doi:10.1093/bioinformatics/btq046)

29. French J P. 2017 autoimage: Multiple Heat Maps for Projected Coordinates. R J. 9, 284. (doi:10.32614/RJ-2017-025)

30. Fox J et al. 2021 car: Companion to Applied Regression. See https://CRAN.R-project.org/package=car.

31. Kassambara A. 2021 rstatix: Pipe-Friendly Framework for Basic Statistical Tests. See https://CRAN.R-project.org/package=rstatix.

32. Landowski M, Kunicka-Kowalska Z, Sibilski K. 2020 Mechanical and structural investigations of wings of selected insect species. Acta Bioeng. Biomech. 22. (doi:10.37190/ABB-01525-2019-03)

33. Racheli T, Pariset L. 1992 Il Genere Battus (Lepidoptera, Papilionidae): Tassonomica e Storia Naturale. See https://www.pemberleybooks.com/product/il-genere-battus-lepidoptera-papilionidae-tassonomica-e-storia-naturale/2784/ (accessed on 13 January 2022).

34. DeVries PJ. 2002 Differential Wing Toughness in Distasteful and Palatable Butterflies: Direct Evidence Supports Unpalatable Theory., 7.

35. Kuznetsova A, Brockhoff PB, Christensen RHB. 2017 lmerTest Package: Tests in Linear Mixed Effects Models. J. Stat. Softw. 82. (doi:10.18637/jss.v082.i13)

36. Naef-Daenzer L, Naef-Daenzer B, Nager RG. 2000 Prey selection and foraging performance of breeding Great Tits Parus major in relation to food availability. J. Avian Biol. 31, 206–214. (doi:10.1034/j.1600-048X.2000.310212.x)

37. Alcock J. 1996 Male size and survival: the effects of male combat and bird predation in Dawson’s burrowing bees, Amegilla dawsoni. Ecol. Entomol. 21, 309–316. (doi:10.1046/j.1365-2311.1996.00007.x)

38. Carvalho MRM, Peixoto PEC, Benson WW. 2016 Territorial clashes in the Neotropical butterfly Actinote pellenea (Acraeinae): do disputes differ when contests get physical? Behav. Ecol. Sociobiol. 70, 199–207. (doi:10.1007/s00265-015-2042-6)

39. Foster DJ, Cartar RV. 2011 What causes wing wear in foraging bumble bees? J. Exp. Biol. 214, 1896–1901. (doi:10.1242/jeb.051730)

40. McKenna KZ, Kudla AM, Nijhout HF. 2020 Anterior–Posterior Patterning in Lepidopteran Wings. Front. Ecol. Evol. 8, 146. (doi:10.3389/fevo.2020.00146)

41. Clarke CA, Sheppard PM, Thornton IWB. 1968 The genetics of the mimetic butterfly Papilio memnon L. Philos. Trans. R. Soc. Lond. B. Biol. Sci. 254, 37–89. (doi:10.1098/rstb.1968.0013)

42. Llaurens V, Whibley A, Joron M. 2017 Genetic architecture and balancing selection: the life and death of differentiated variants. Mol. Ecol. 26, 2430–2448. (doi:10.1111/mec.14051)

43. Rawlins JE. 1980 Thermoregulation by the Black Swallowtail Butterfly, Papilio Polyxenes (Lepidoptera: Papilionidae). Ecology 61, 345–357. (doi:10.2307/1935193)

44. Kim Y, Hwang Y, Bae S, Sherratt TN, An J, Choi S-W, Miller JC, Kang C. 2020 Prey with hidden colour defences benefit from their similarity to aposematic signals. Proc. R. Soc. B Biol. Sci. 287, 20201894. (doi:10.1098/rspb.2020.1894)

45. Holmes GG, Delferrière E, Rowe C, Troscianko J, Skelhorn J. 2018 Testing the feasibility of the startle-first route to deimatism. Sci. Rep. 8, 10737. (doi:10.1038/s41598-018-28565-w)

46. Dudley R. 2002 The Biomechanics of Insect Flight: Form, Function, Evolution. Princeton University Press.

47. Park H, Bae K, Lee B, Jeon W-P, Choi H. 2010 Aerodynamic Performance of a Gliding Swallowtail Butterfly Wing Model. Exp. Mech. 50, 1313–1321. (doi:10.1007/s11340-009-9330-x)

48. Basset J-P, Cartraud P, Jacquot C, Leroy A, Peseux B, Vaussy P. In press. Introduction à la résistance des matériaux., 179.

