## Supplementary materials S1: Physical characterisation of the tensile strength of the wing (from [48]). for "Evidence of attack deflection suggests adaptive evolution of wing tails in butterflies": Supplementary S1.docx

**Physical characterisation of the tensile strength of the wing (Basset et al 2007)**

The behaviour of a material in a tensile test is given by Hooke's law:

[1]

$$\sigma=E.\varepsilon$$

With: 𝐸 the Young's modulus of the material (an intrinsic modulus of elasticity of the material)

𝜎 the stress applied to the sample (N/mm2)

𝜀 the resulting strain of the specimen

The stress and strain of the sample are given by the following equations

$$\sigma= \frac{F}{S}$$

[2]

ε $= \frac{L-Lo}{Lo}$

[3]

With: F the force applied to the sample (N)

S the area of the sample normal to the force (mm^2^)

L_o_ the initial length of the sample before traction (mm)

L the length of the sample after traction (mm)

Thus, according to the equation [1] the behaviour of a material in a tensile test can be characterised by the following equation:

$$F=E.S.\frac{L-Lo}{Lo}$$

[4]

Knowing that our results represent the force exerted on a vein as a function of time, and that the elapsed time is proportional to the deformation of the material (because the tensile force is applied to the wing in a constant manner), then the slope measured for each curve profile is equivalent to:

[5]

$$\beta=E.S$$

The slope measured on our curve gives us an idea of the order of magnitude of the Young's modulus. The greater the slope, the stiffer the vein.
