## Supplementary figures and images for "Evidence of attack deflection suggests adaptive evolution of wing tails in butterflies"

### Supplementary S2.jpg

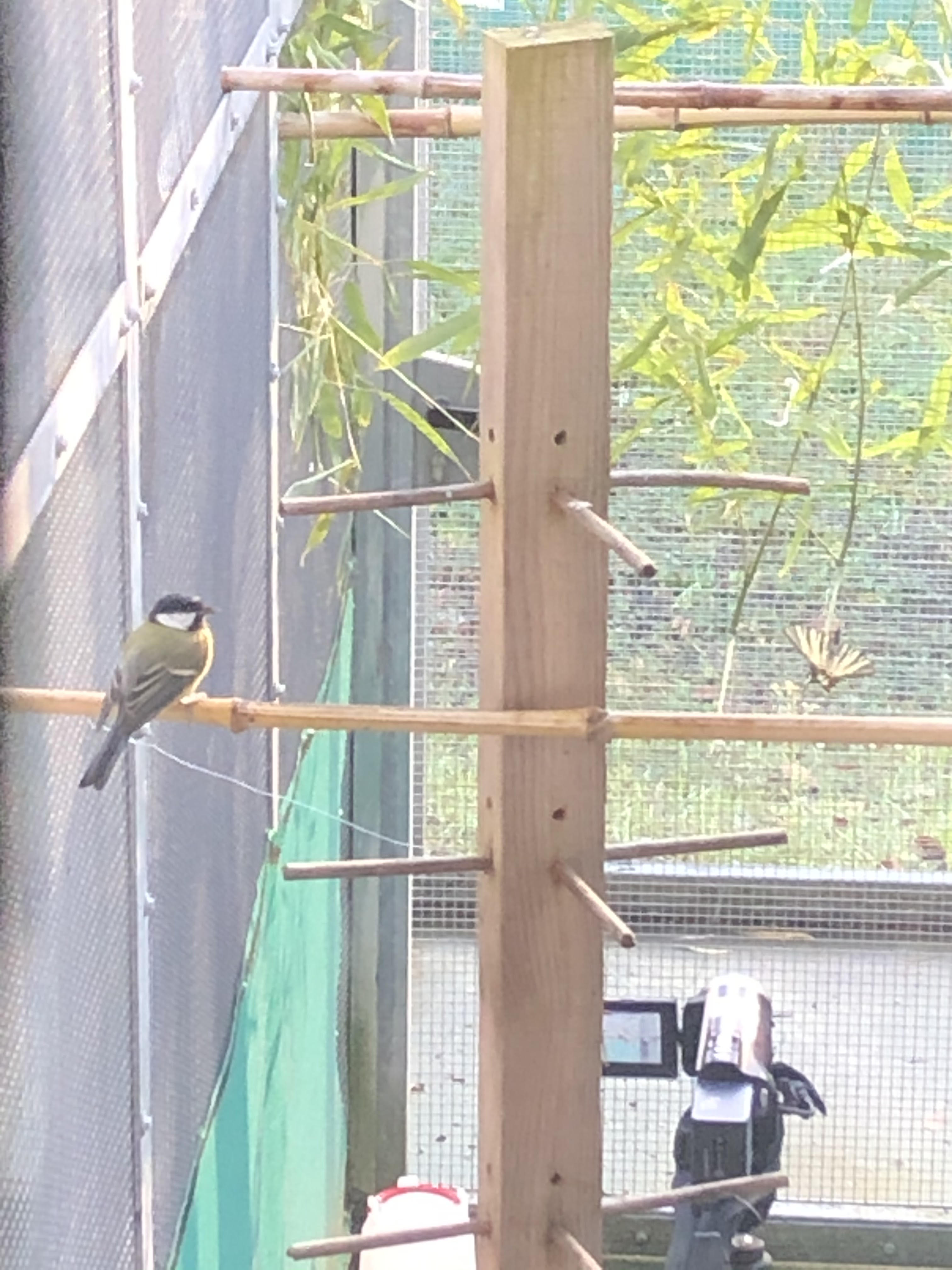

### Supplementary S3.JPG

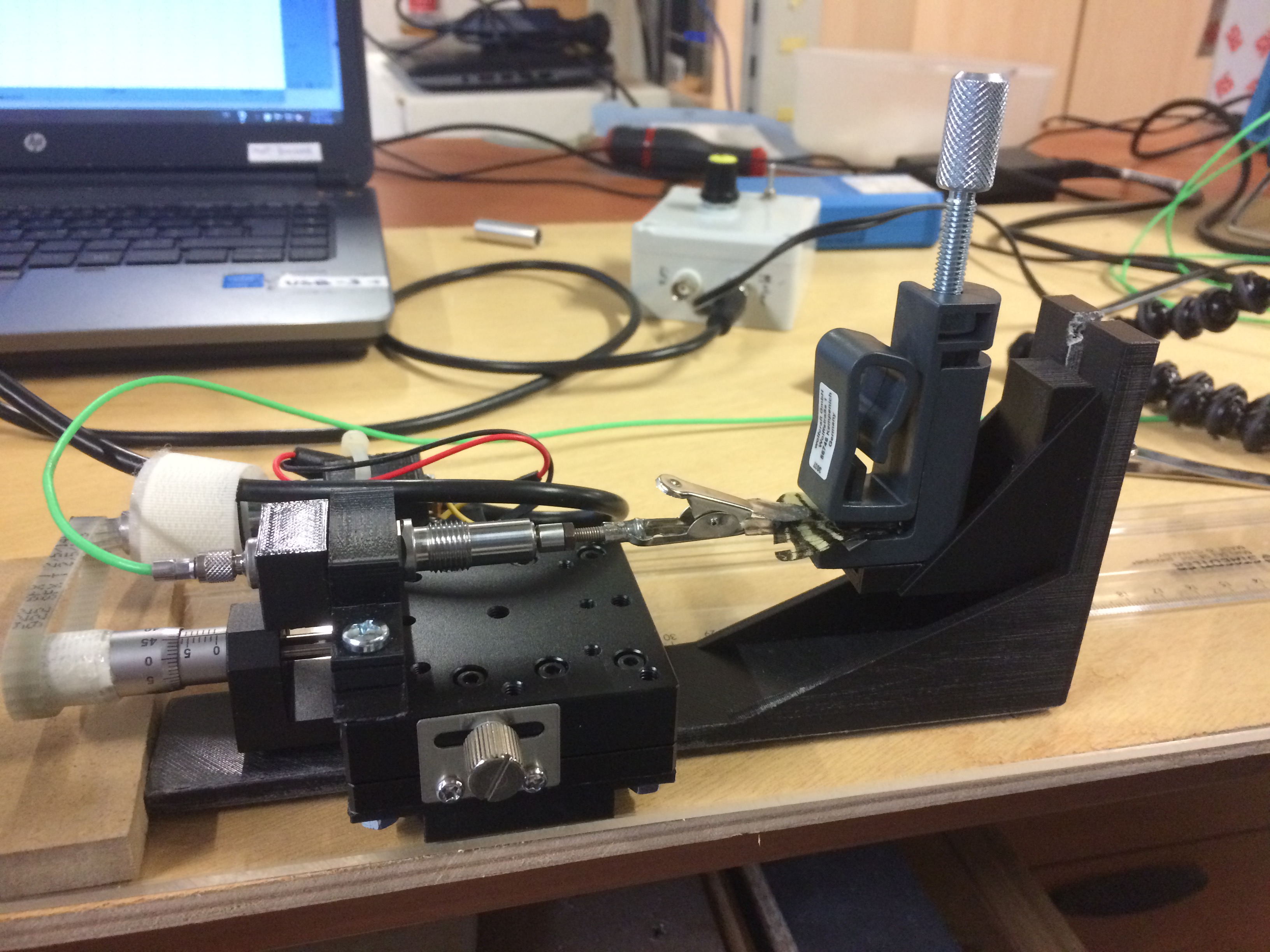

### Supplementary S4.pdf

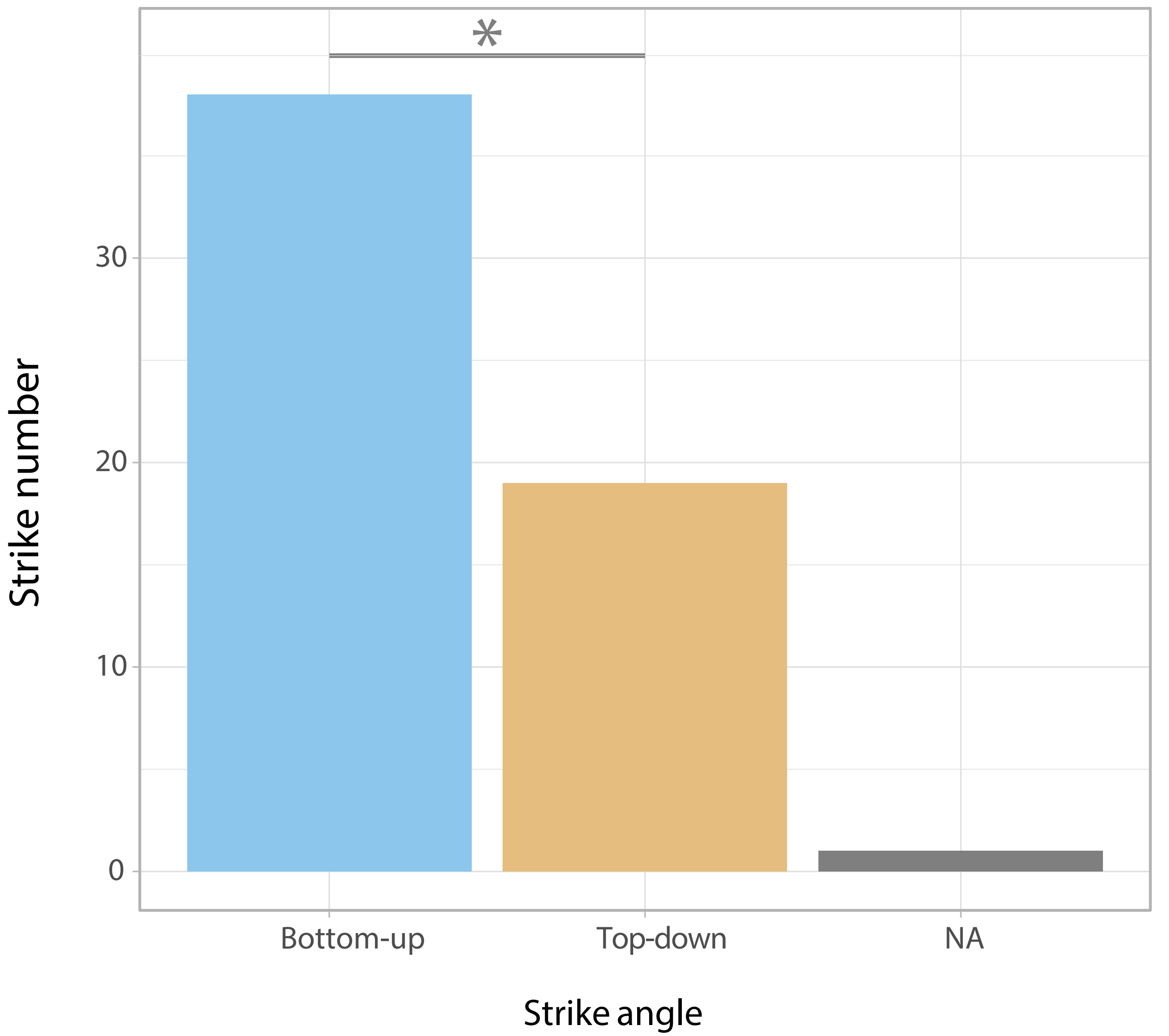
